# Genomic signals of admixture and reinforcement between two closely related species of European sepsid flies

**DOI:** 10.1101/2020.03.11.985903

**Authors:** Athene Giesen, Wolf U. Blanckenhorn, Martin A. Schäfer, Kentaro K. Shimizu, Rie Shimizu-Inatsugi, Bernhard Misof, Oliver Niehuis, Lars Podsiadlowski, Heidi E. L. Lischer, Simon Aeschbacher, Martin Kapun

## Abstract

Interspecific gene flow by hybridization may weaken species barriers and adaptive divergence, but can also initiate reinforcement of reproductive isolation trough natural and sexual selection. The extent of interspecific gene flow and its consequences for the initiation and maintenance of species barriers in natural systems remain poorly understood, however. To assess genome-wide patterns of gene flow between the two closely related European dung fly species *Sepsis cynipsea* and *Sepsis neocynipsea* (Diptera: Sepsidae), we tested for historical gene flow with the aid of ABBA-BABA test using whole-genome resequencing data from pooled DNA of male specimens originating from natural and laboratory populations. We contrasted genome-wide variation in DNA sequence differences between samples from sympatric populations of the two species in France and Switzerland with that of interspecific differences between pairs of samples involving allopatric populations from Estonia and Italy. In the French Cevennes, we detected a relative excess of DNA sequence identity, suggesting interspecific gene flow in sympatry. In contrast, at two sites in Switzerland, we observed a relative depletion of DNA sequence identity compatible with reinforcement of species boundaries in sympatry. Our results suggest that the species boundaries between *S. cynipsea* and *S. neocynipsea* in Europe depend on the eco-geographic context.

## INTRODUCTION

According to the biological species concept, species represent groups of individuals that are reproductively isolated from other groups of individuals (Mayr 1942). Speciation entails the evolution of reproductive isolation among lineages derived from a common ancestral population and is considered completed if the diverged populations remain reproductively isolated, for example, after coming into secondary contact (Coyne & Orr, 2004; Dobzhansky, 1951; Mayr, 1963). However, many animal and plant species remain distinct entities in nature even if they occasionally hybridize and exchange genes in parapatry or sympatry (Anderson, 1949; Barton & Bengtsson, 1986; cf. DeMarais *et al*., 1992; Gante *et al*., 2016; Mallet, 2007; Nolte & Tautz, 2010; Rieseberg *et al*., 2003; Trier *et al*., 2014). Research during the past decades has indicated that hybridization not only has deleterious effects due to hybrid inferiority and negative epistasis in admixed genomes, but that it may also fuel adaptive diversification and speciation by facilitating novel combinations of alleles that become targets of divergent selection (Arnold & Meyer, 2006; Berner & Salzburger, 2015; Fontaine *et al*. 2015; Seehausen, 2004; Saetre, 2013). Together, these findings corroborate the view that genomes of recently diverged species may represent mosaics, consisting of genomic regions with significant differentiation (i.e., low to no admixture) interspersed by genomic regions that are more free to be exchanged (Nosil *et al*., 2009; Wu, 2001).

Sepsid flies (Diptera: Sepsidae) have become a model group for the study of sexual selection and ecological adaptation (Baur *et al*., 2020; Blanckenhorn, 1999; Blanckenhorn *et al*., 2000; Eberhard, 1999; Kraushaar & Blanckenhorn, 2002; Parker, 1972a,b; Puniamoorthy *et al*., 2009; Pont & Meier, 2002; Rohner *et al*., 2015; Rohner, Blanckenhorn, & Puniamoorthy, 2016; Ward, 1983; Ward, Hemmi, & Rösli, 1992). The phylogeny of sepsid flies is well resolved (Su *et al*., 2008, 2016) and entails multiple pairs of closely related species that occupy similar ecological niches. These species pairs provide excellent opportunities to study the genomic consequences of hybridization and introgression during early stages of speciation. One of these pairs comprises *Sepsis cynipsea* and *Sepsis neocynipsea*, sister species with a wide geographic distribution that occur in sympatry across major parts of their natural range. While *S. cynipsea* is the most abundant sepsid species in Central and Northern Europe and deposits its eggs into fresh cow dung, *S. neocynipsea* is common throughout North America, where it occupies a niche similar to that of *S. cynipsea* in Europe. While overall very rare in Europe, *S. neocynipsea* can be locally common at higher altitudes, such as the Alps, where the species occurs in sympatry with *S. cynipsea* (Ozerov, 2005; Pont & Meier, 2002; Rohner, Blanckenhorn, & Puniamoorthy, 2016). Despite strong similarities in morphology and behavior (Giesen, Blanckenhorn, & Schäfer, 2017), the two species are genetically distinct. Previous studies showed that *S. cynipsea* and *S. neocynipsea* produce fertile hybrid offspring under laboratory conditions despite strong pre- and post-mating isolating barriers. These barriers are mediated by assortative mating behaviors that are partly reinforced in areas where the two species occur in sympatry (Giesen *et al*., 2017, 2019). While these findings imply that interspecific gene flow may occur and vary with eco-geographic context in nature, we know little about actual levels of historical and ongoing gene flow between the two species in areas where they occur in sympatry or parapatry. We therefore investigated the extent of gene flow between *S. cynipsea* and *S. neocynipsea* in nature by comparative population genomic analyses.

In recent years, multiple approaches have been employed to explore the importance of interspecific gene flow in sympatry vs. allopatry (e.g. Nakazato, Warren, & Moyle, 2010; Nadeau *et al*., 2013; Brandvain *et al*., 2014; Bouchemousse *et al*., 2016; Feulner & Seehausen, 2019; Kastally, Trasoletti, & Mardulyn, 2019; Moran *et al*., 2019). The objective of our study is to assess the extent of genome-wide admixture between *S. cynipsea* and *S. neocynipsea* in Europe by applying a version of the ABBA-BABA test for historical gene flow (Green *et al*., 2010; Durand *et al*., 2011; Soraggi et al., 2018) that can exploit genome-wide allele frequency data of single nucleotide polymorphisms (SNPs). To this end, we sequenced the pooled genomic DNA of males from wild-caught populations of *S. cynipsea* and *S. neocynipsea* collected at multiple sites ranging from Southern France to Estonia. Some of these populations occur in sympatry (e.g., in the Swiss Alps and the French Cevennes), others in allopatry (Pont & Meier, 2002). Based on hybridization opportunity alone, we expected to find higher genome-wide levels of gene flow between *S. cynipsea* and *S. neocynipsea* in geographic areas where both species occur in sympatry (e.g. Nadeau *et al*., 2013, or Martin *et al*., 2014, for *Heliconius* butterflies). In contrast, less gene flow in sympatry than in allopatry would indicate selection against gene flow and thus suggest reinforcement of reproductive barriers at sites of co-occurrence (Butlin, 1995; Noor, 1999; Coyne & Orr, 2004; e.g. Kulathinal & Singh, 2000; Massie & Makow, 2005; Giesen *et al*. 2017, 2019).

## MATERIALS & METHODS

### Sample collection and treatment of fly cultures

We studied inter- and intraspecific gene flow in S. *cynipsea* and *S. neocynipsea* using genomic data of sympatric *S. cynipsea* and *S. neocynipsea* populations from two high altitude sampling sites in the Swiss Alps (Sörenberg) and the French Cevennes (Le Mourier) (Table 1). To complement these two pairs of sympatric populations with geographically distant allopatric populations in Europe (Table 1), we further included *S. cynipsea* samples from the Swiss lowlands (Zürich), from Tuscany, Italy (Petroia) and Estonia (Pehka), and from two additional high-altitude *S. neocynipsea* samples collected in Switzerland (Geschinen and Hospental) (Table 1). In line with the previous observation that *S. neocynipsea* tends to be rare at low altitudes (Pont & Meier, 2002), we were not able to collect sufficient numbers of flies of this species outside the Alps, except for the sample collected near Le Mourier. For each of the wild-caught populations, we randomly selected 20 males for pooled DNA resequencing (Pool-Seq).

**Table 1:**
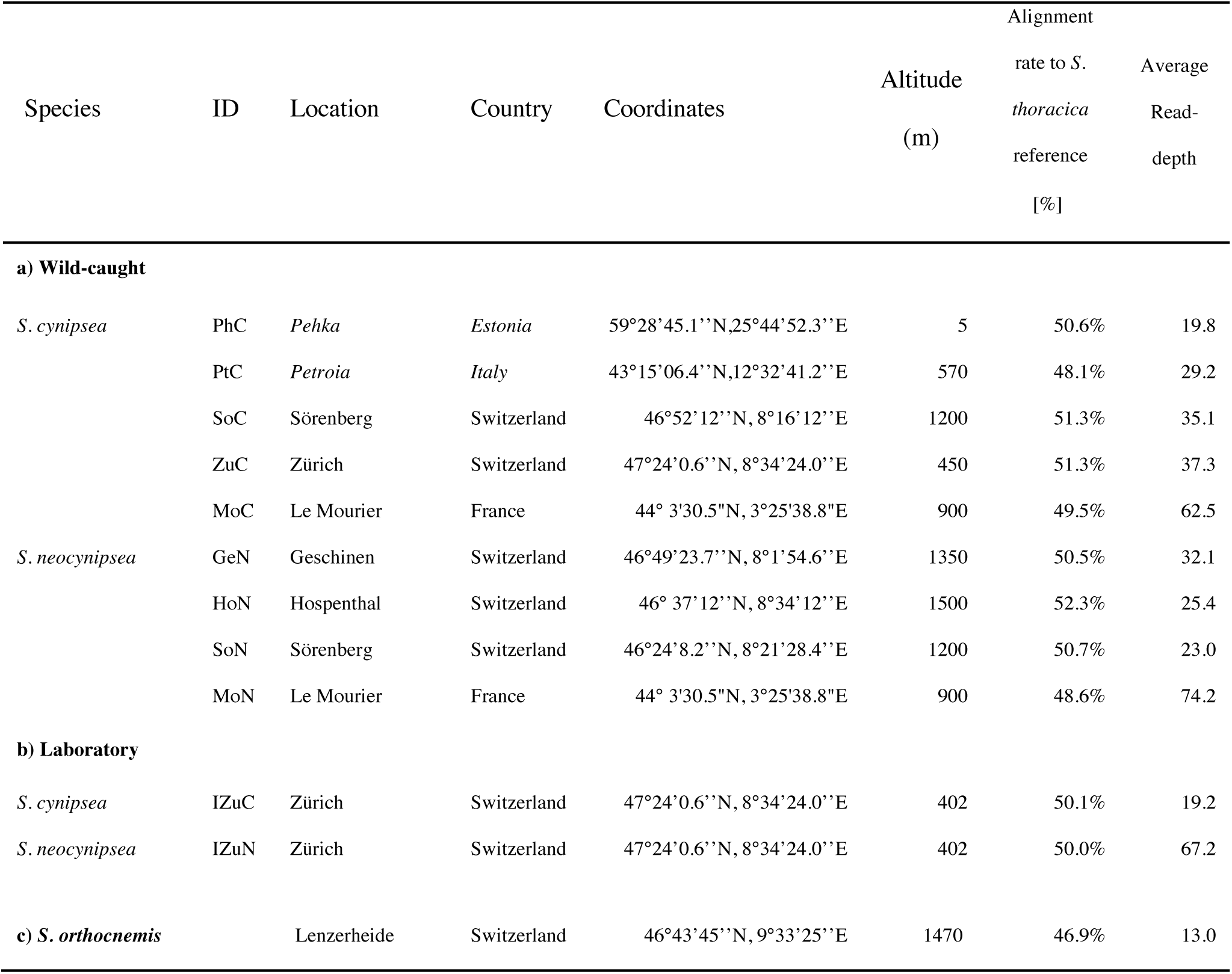
Sampling sites, sample IDs, percentages of DNA sequence reads mapping to the reference genome and read-depths of pool-sequenced field-collected (a) and laboratory (b) populations. Sites where *S. cynipsea* and *S. neocynipsea* occur in sympatry are in regular font. Sites where *S. neocynipsea* does not occur to our knowledge are highlighted in italics.

| Species | ID | Location | Country | Coordinates | Altitude<br>(m) | Alignment<br>rate to *S.*<br>*thoracica*<br>reference<br>[%] | Average<br>Read-<br>depth |
| --- | --- | --- | --- | --- | --- | --- | --- |
| **a) Wild-caught** |  |  |  |  |  |  |  |
| *S. cynipsea* | PhC | *Pehka* | *Estonia* | 59°28'45.1''N, 25°44'52.3''E | 5 | 50.6% | 19.8 |
|  | PtC | *Petroia* | *Italy* | 43°15'06.4''N, 12°32'41.2''E | 570 | 48.1% | 29.2 |
|  | SoC | Sörenberg | Switzerland | 46°52'12''N, 8°16'12''E | 1200 | 51.3% | 35.1 |
|  | ZuC | Zürich | Switzerland | 47°24'0.6''N, 8°34'24.0''E | 450 | 51.3% | 37.3 |
|  | MoC | Le Mourier | France | 44° 3'30.5"N, 3°25'38.8"E | 900 | 49.5% | 62.5 |
| *S. neocynipsea* | GeN | Geschinen | Switzerland | 46°49'23.7''N, 8°1'54.6''E | 1350 | 50.5% | 32.1 |
|  | HoN | Hospenthal | Switzerland | 46° 37'12''N, 8°34'12''E | 1500 | 52.3% | 25.4 |
|  | SoN | Sörenberg | Switzerland | 46°24'8.2''N, 8°21'28.4''E | 1200 | 50.7% | 23.0 |
|  | MoN | Le Mourier | France | 44° 3'30.5"N, 3°25'38.8"E | 900 | 48.6% | 74.2 |
| **b) Laboratory** |  |  |  |  |  |  |  |
| *S. cynipsea* | IZuC | Zürich | Switzerland | 47°24'0.6''N, 8°34'24.0''E | 402 | 50.1% | 19.2 |
| *S. neocynipsea* | IZuN | Zürich | Switzerland | 47°24'0.6''N, 8°34'24.0''E | 402 | 50.0% | 67.2 |
| **c) *S. orthocnemis*** |  | Lenzerheide | Switzerland | 46°43'45''N, 9°33'25''E | 1470 | 46.9% | 13.0 |

In addition to wild-caught populations, we compiled genomic data from two laboratory populations that were established from sympatric *S. cynipsea* and *S. neocynipsea* populations collected near Zürich. We started each of these populations from a single gravid, wild-caught female and subsequently propagated them in the laboratory for at least two years at census population sizes ranging from 10 to 100. We then sampled and pooled the DNA of 50 males from each population for whole-genome sequencing (see Supporting Information for more details).

### DNA library preparation and next generation sequencing

Genomic DNA was extracted from the pooled DNA of males using the UltraPure Phenol:Chloroform:Isoamyl alcohol (25:24:1, v/v) extraction kit (Thermo Fischer Scientific, Waltham, USA) according to the manufacturer’s protocol. Quantification of genomic DNA was performed with a Qubit Fluorometer (Thermo Fischer Scientific; Table 1). Library preparation was carried out with the TruSeq DNA PCR-Free Library Preparation (Illumina, San Diego, USA) kit according to the manufacturer’s protocol. Fragment-size distributions of all libraries were validated on a TapeStation 2200 (Agilent Technologies, Waldbronn, Germany). Sequencing on Illumina HiSeq 2500 version 4 was conducted after labeling and pooling the barcoded DNA representing the four laboratory populations onto one lane to achieve a genome-wide coverage of ca. 60x per DNA pool library. All sequencing data are available at the short-read archive (SRA; https://www.ncbi.nlm.nih.gov/sra) under the accession number PRJNA612154.

### Raw read processing

Qualitative validation of sequence data before and after trimming was done with FastQC (v. 0.11.4; Andrews *et al*., 2011). After removal of Illumina-specific adapters and trimming with *Trimmomatic* (v. 0.36; Bolger, Lohse & Usadel, 2014), the *S. cynipsea* and *S. neocynipsea* reads were mapped to the draft genome of *S. thoracica* with *bwa mem* (v. 0.7.12; Li & Durbin, 2009) using default parameters. The *S. thoracica* sample used for genome sequencing was collected near Capriasca, Ticino, Switzerland. The draft genome, v. 0.1 was built from Oxford Nanopore long reads (app. 25x read depth), assembled with Canu (Korner *et al*., 2017), and polished with Illumina short reads (app. 20x read depth). The assembly used in this study is available under the rules of the Fort Lauderdale agreement from www.cgae.de/seto_01_genome.fasta and www.cgae.de/seto_01_genome_masked.fasta.

PCR duplicates were removed with Picard (v. 1.109; http://broadinstitute.github.io/picard/) and reads were realigned around indels in each raw alignment file using RealignerTargetCreator and IndelRealigner from GATK (v. 3.4-46; McKenna et al. 2010). Only reads with a PHRED-scaled mapping quality of 20 and more were retained. We applied the same procedure to map reads of a pool of ten inbred *S. orthocnemis* males from near Lenzerheide, Switzerland, to the reference genome (Table 1). We compiled the aligned reads from all population samples into a single *mpileup* file using *samtools* (v. 1.3.1; Li *et al*., 2009).

Using the variant-caller Pool-SNP (Kapun *et al*., 2018), we identified high-confidence SNPs with the following combination of heuristic SNP-calling parameters: coverage of each Pool-Seq sample ≥ 10x; coverage of each Pool-Seq sample ≤ 95 % percentile of coverage distribution across contigs and samples; minimum allele count of a minor allele at a SNP across all combined Pool-Seq samples > 20x; minor allele frequency at a SNP across all combined Pool-Seq samples > 0.01. We only retained SNPs for which all samples fulfilled the above coverage threshold criteria. To avoid paralogous SNPs due to mis-mapping around indel polymorphisms, we removed SNPs located within a 5-bp distance to indels that occurred in more than 20 copies at a given site across all pooled samples. In addition, we ignored SNPs located within repetitive regions. The resulting VCF file was converted to the SYNC file format (Kofler *et al*., 2011), and a custom Python script was used to calculate sample-specific allele frequencies for major alleles at each SNP (*sync2AF*.*py*; https://github.com/capoony/DrosEU_pipeline).

### Testing for introgression

To test for introgression, we used the ABBA-BABA test for historical gene flow in the presence of incomplete lineage sorting (Green *et al*., 2010; Durand *et al*., 2011). We provide a brief motivation of the approach in the following and refer to the Supporting Information for further details. Imagine a rooted phylogeny with four species (P1 to P4) and a topology of (((P1, P2), P3), P4) as illustrated in Fig. 1B. In an alignment of one haploid genome from each species, incomplete lineage sorting under neutral evolution leads to two mutational configurations of the ancestral (A) and derived (B) allele, ABBA and BABA, that are incompatible with the species topology (Fig. 1B) under the infinite-sites model of mutation (Kimura, 1969). In the absence of gene flow between P2 and P3, ABBA and BABA configurations occur with equal probability (Hudson, 1983; Tajima, 1983). In contrast, historical gene flow between P2 and P3 causes an excess of ABBA configurations (Green *et al*., 2010; Durand *et al*., 2011). This logic is captured by the *D*-statistic, a scaled difference across a set of bases between counts of ABBA and BABA configurations bounded by –1 and 1 (Green *et al*., 2010). The ABBA-BABA test examines significant deviations of *D* from 0. A significant excess of ABBA suggests evidence for gene flow between P2 and P3. A significant depletion of ABBA is equivalent to an excess of ABBA if the positions of P1 and P2 in the species topology are swapped, and hence either suggests gene flow between P1 and P3, or a reduction in gene flow between P2 and P3 relative to gene flow between P1 and P3.

**Fig. 1.**
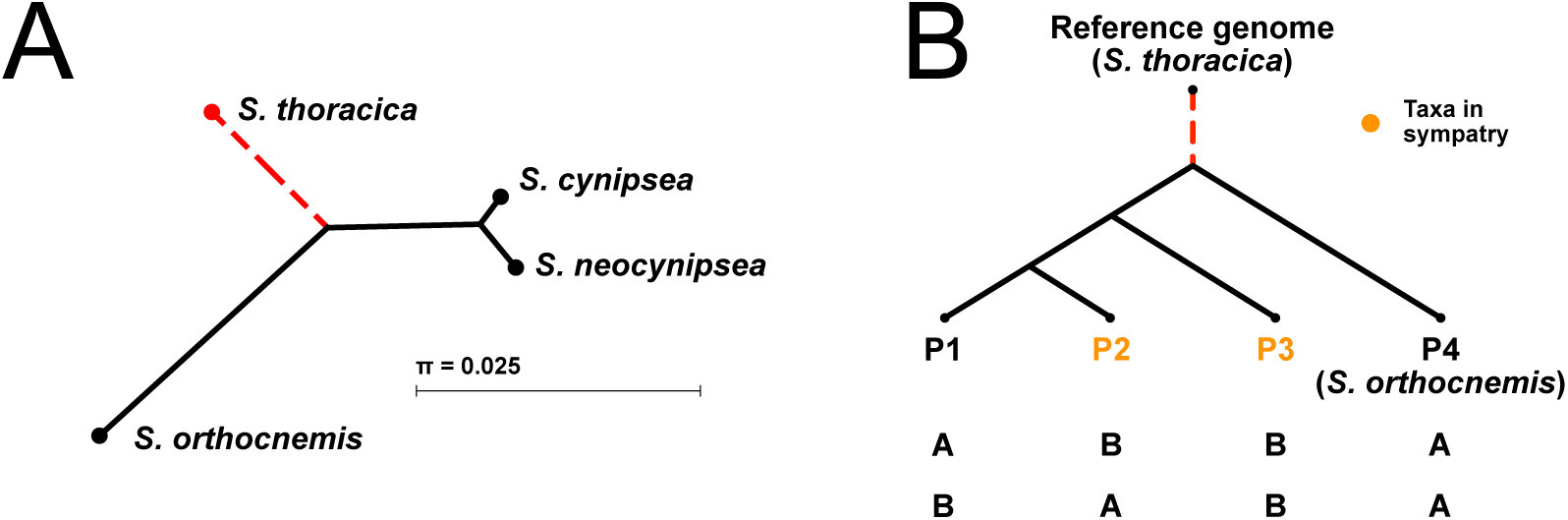
Putative phylogeny of sepsid species studied. (A) Species phylogeny inferred from alignments of nucleotide sequences at the cytochrome oxidase subunit II (CO-II) mitochondrial locus (Genbank sequences from Su *et al*., 2016) with the Neighbor-Joining method implemented in CLC Main Workbench (v. 8.1.2; https://www.qiagenbioinformatics.com/products/clc-main-workbench/). The tree topology is consistent with the phylogeny presented by Su *et al*.*’s* (2016) combined phylogenetic analysis of multiple nuclear and mitochondrial markers. *Sepsis orthocnemis* was used as the outgroup for the focal species *S. cynipsea* and *S. neocynipsea*. Whole-genome sequencing reads from all three species were aligned to the *S. thoracica* reference genome (dashed red branches). Branch lengths are proportional to the mean number of pairwise sequence differences. (B) Generic species tree assumed in all our ABBA-BABA tests for gene flow among triplets of *S. cynipsea* and *S. neocynipsea* populations from various sites in Europe.

The original version of the ABBA-BABA test and modifications of it have been applied to different types of genome-scale polymorphism data obtained with various sequencing strategies, including whole genome sequencing (e.g. Green *et al*., 2010), RAD sequencing (e.g., Eaton & Ree, 2013; Meier *et al*., 2017; Streicher *et al*., 2014), and exon capture data (e.g., Heliconius Genome Consortium, 2012). More recently, Durand *et al*. (2011) and Soraggi *et al*. (2018) extended the original test to allele frequency data, hence to unphased sequencing data (including Pool-Seq data; Schlötterer *et al*. 2014). We implemented the extensions by Durand *et al*. (2011) and Soraggi *et al*. (2018) in a Python script (https://github.com/capoony/ABBABABA-4AF) and refer to the respective test statistics as *D*_D_ and *D*_S_. Our script also computes jack-knifed *z*-scores based on a matrix of allele frequencies for previously defined high-confidence SNPs (see Supporting Information for details). We adopted the commonly used significance threshold of |*z*| > 3 (Reich *et al*., 2011; Jeong *et al*., 2016; Novikova *et al*., 2016) to identify significant deviations of *D*_S_ and *D*_D_ from zero.

We were concerned that using the same genome assembly both as the reference for read mapping and as the outgroup (P4) in the ABBA-BABA tests could lead to biased estimates of gene flow. We therefore mapped the three ingroup and the outgroup species against the reference genome of yet another species, *S. thoracica*. A phylogenetic analysis based on CO-II sequences by Su *et al*. (2008) indicated that *S. thoracica* is approximately equally related to *S. cynipsea* and *S. neocynipsea* as it is from the outgroup species *S. orthocnemis* (Fig. 1A). We therefore expected that reads of the ingroup and the outgroup species would map equally well to the reference species, which is indeed what we observed (Table 1).

We focused our analysis of historical interspecific gene flow on the three sampling sites Sörenberg, Zürich and Le Mourier (Table 1). At all these sites, *S. cynipsea* and *S. neocynipsea* occur in sympatry. In all ABBA-BABA tests, we positioned the two focal populations as P2 and P3 ingroups in the phylogeny (in both possible orders) and used various P1 ingroup populations (Fig. 1B) assumed to occur in allopatry with the P2 populations. We used various P1 ingroups, because Durand *et al*. (2011) showed that the choice of the ingroup P1 can influence the test results.

## RESULTS

### Signals of reinforcement in Swiss alpine and sub-alpine regions

The majority of our tests for gene flow at the Sörenberg site revealed a genome-wide deficiency of derived-allele sharing (i.e., ABBA patterns) among the local *S. cynipsea* and *S. neocynipsea* populations (Table 2), suggesting that interspecific gene flow in sympatry (among P2 and P3) is lower than interspecific gene flow among putatively allopatric populations (P1 and P3). This observation is robust to our choice of the P1 ingroup population, but sensitive to the version of *D*-statistic used (*D*_S_ vs. *D*_D_). We first conducted tests involving two *S. neocynipsea* populations from the Swiss Alps (*S. neocynipsea* from Geschinen, GeN; *S. neocynipsea* from Hospental, HoN; Table 1) as putatively allopatric P1, and the sympatric interspecific pair from Sörenberg as P2 (*S. neocynipsea*) and P3 (*S. cynipsea*). We observed a significant deficiency of ABBA patterns in all configurations (*D*_S_ > 0; Table 2, upper part), suggesting overall evidence for higher gene flow among putatively allopatric than sympatric interspecific population pairs. While GeN and HoN are both separated by high mountains (altitudes ranging from ca. 600–3,200 m asl) from the Sörenberg site, they are geographically close (ca. 67.5 km and ca. 46.7 km Euclidean distance for GeN and HoN, respectively) and thy occur in sympatry with other sepsid species, including *S. cynipsea*. We were therefore concerned that our assumption of no gene flow between P1 and P3 was violated, because GeN and HoN are parapatric rather than allopatric to our focal populations in Sörenberg.

**Table 2:** Results of ABBA-BABA tests applied to samples originating from sympatric populations of *S. cynipsea* and *S. neocynipsea* in Sörenberg as P2 and P3 (or vice versa) based on *D*_S_. The upper half of the table shows tests with two European *S. neocynipsea* populations from Geschinen (GeN) and Hospental (HoN) as P1 ingroups, and the bottom half shows the reciprocal approach with two *S. cynipsea* populations from Pehka, Estonia (PhC) and Petroia, Italy (PtC) as P1 ingroups. The first three columns indicate the phylogeny assumed for the test, with P4 always being *S. orthocnemis*; columns 4 to 6 give the genome-wide average *D*_S_, its jackknifed standard deviation SD(*D*_S_), and the corresponding *z*-score. Significant tests are shown in bold.

| P1 | P2 | P3 | $D_S$ | $SD(D_S)$ | $z$ |
| --- | --- | --- | --- | --- | --- |
| **GeN** | **SoN** | **SoC** | **0.008** | **1.35E-03** | **5.8** |
| **HoN** | **SoN** | **SoC** | **0.006** | **1.34E-03** | **4.2** |
| **PhC** | **SoC** | **SoN** | **0.026** | **1.92E-03** | **13.4** |
| **PtC** | **SoC** | **SoN** | **0.039** | **1.97E-03** | **19.8** |

To address this concern, we conducted additional ABBA-BABA tests involving P1 populations from geographically more remote sites in Europe. Specifically, we performed a second series of tests with two alternative *S. cynipsea* populations as allopatric P1 ingroups (Fig. 1B), one from Pehka, Estonia (PhC), and one from Petroia, Italy (PtC; Table 1). Contrary to our expectation based on the much larger geographical distance between P1 and P3 (ca. 1,850 km and ca. 550 km for PhC and PtC, respectively) than between P2 and P3 in these configurations, we still found a significant deficiency of ABBA patterns in both tests (*D*_S_ > 0; Table 2, lower part). In agreement with our previous tests involving GeN and HoN as P1, these additional tests thus equally suggest higher interspecific gene flow among allopatric (P1 and P3) than among sympatric populations (P2 and P3). As we are much more confident that PhC and PtC represent truly allopatric P1 ingroups relative to P2 and P3, we think that the deficiency of ABBA patterns at the Sörenberg site indicates a local reduction in interspecific gene flow in sympatry, likely due to reinforcement.

When repeating the ABBA-BABA tests with the *D*-statistic (*D*_D_) suggested by Durand *et al*. (2011), we obtained results that qualitatively agree with those obtained with the *D*_S_ statistic only when using PhC and PtC as P1 (*D*_D_ < 0; z scores -8.4 and -7.7, respectively; Table S1 lower part), but not with GeN and HoN as P1 (*z*-scores 1.4 and 1.9, respectively; Table S1, upper part). Besides the above qualitative differences between tests based on *D*_S_ vs. *D*_D_, we observed that variances of *D*_D_ estimated by jackknifing were about one order of magnitude larger than the corresponding variances of *D*_S_, which might indicate differences in the robustness of the two measures. Nonetheless, the qualitatively similar results obtained with the two versions of *D*-statistics makes us confident that our findings are robust. We further observed that windows of elevated *D*_S_ (Fig. 2, top) were not homogeneously distributed along the genome, but aggregated in clusters in several contigs. Such clustering might suggest that the respective genomic regions contain variation involved in reinforcement caused by divergent selection against gene flow between the two species.

**Fig. 2.**
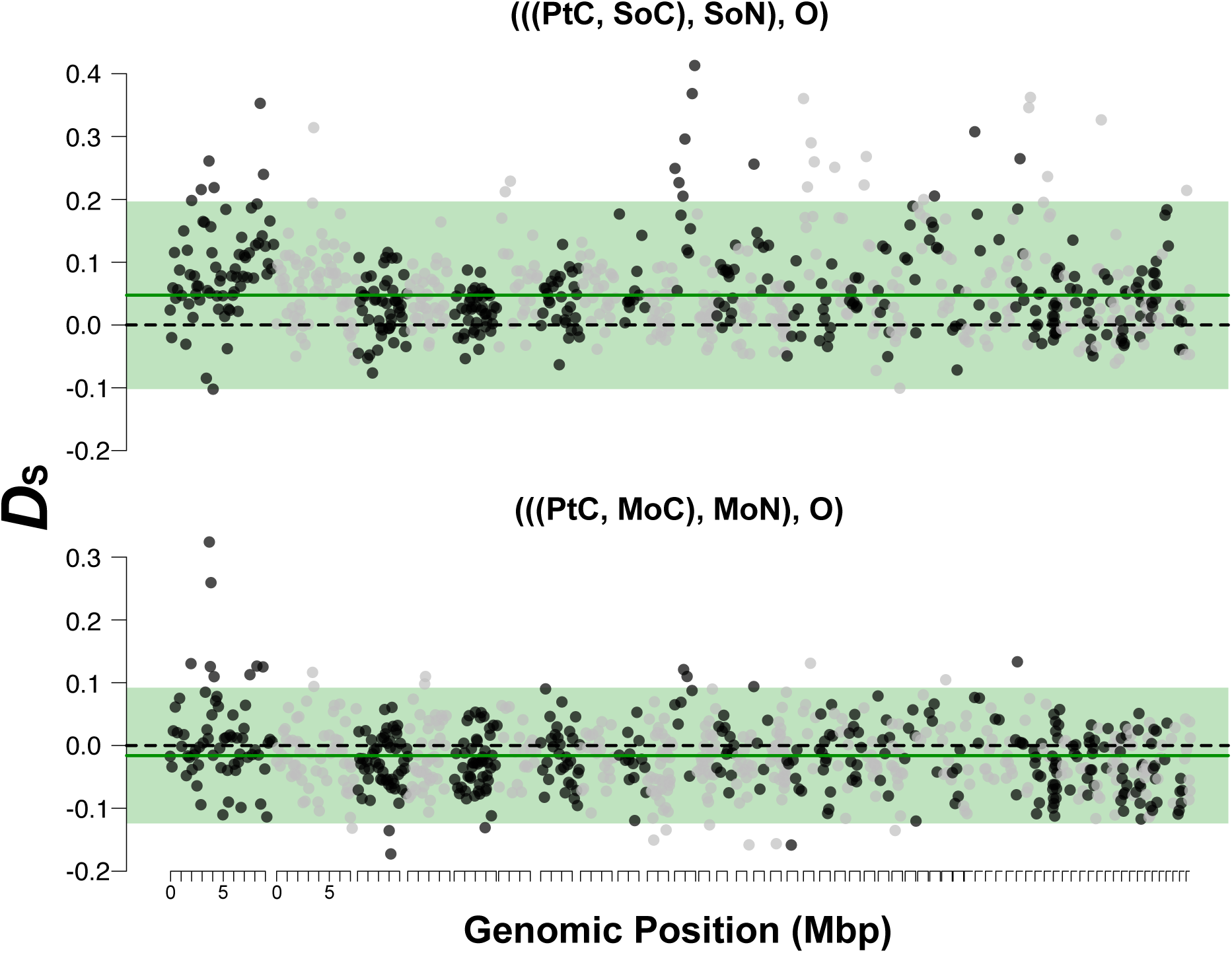
Variation of *D*_S_ along the genome. Dots indicate *D*_S_ (Soraggi *et al*. 2018) for genomic windows of 500 consecutive SNPs, oriented by contigs of the *S. thoracica* reference genome with a length ≥ 50,000 bp (highlighted by alternating black and grey colors). The top and bottom panels show results for the population tree topologies used to test for interspecific gene flow at the focal sites in Sörenberg (SoC-SoN; deficiency of ABBA patterns) and Le Mourier (MoC-MoN; excess of ABBA patterns), respectively. The horizontal dashed black and green lines indicate the neutral expectation (*D*_S_ = 0) and the mean genome-wide jackknife estimate, respectively. Green shading delimits mean *D*_S_ ± 2 standard deviations (based on windows of 500 SNPs).

Since both species commonly co-occur in alpine and sub-alpine regions of Switzerland, we were curious to see if a signal of reinforcement was also evident at other sites of sympatry in this geographic area. We therefore investigated a nearby lowland sampling site in Zürich where we also found *S. cynipsea* and *S. neocynipsea* in sympatry. In contrast to the sequencing data from Sörenberg, which were generated from wild-caught specimens only, the sequencing data from the Zürich site derive from a combination of samples from natural and laboratory populations. Specifically, for *S. cynipsea*, we sequenced a pool of flies sampled directly from the natural population as well as a pool of flies from a laboratory population that was derived from the same natural population (see Material and Methods for details); for *S. neocynipsea*, we only had a pool of flies sampled from a laboratory population derived from the natural population (Table 1). We found that the ABBA-BABA tests were qualitatively robust to whether we used *S. cynipsea* data from natural or laboratory populations. However, results depended on the choice of the *D* statistic (*D*_S_ vs. *D*_D_) and of the P1 ingroup (Supplementary Information, Supplementary Table S2). All ABBA-BABA tests based on *D*_S_ that included GeN, HoN, or PtC as P1 were significant and revealed a deficiency of ABBA patterns, whereas tests with PhC as P1 were not significant (Supplementary Table S2). All significant tests were therefore consistent with our tests for the Sörenberg site in that they suggested reduced interspecific gene flow in sympatry relative to allopatry. In contrast, only two out of eight ABBA-BABA tests based on *D*_S_ were significant, one suggesting an excess and one a deficiency of ABBA patterns. Overall, our results for the Zurich site are compatible with reinforcement, although the signal seems to be weaker compared to the Sörenberg site.

### Signals of introgression in high altitude populations from Southern France

Complementary to the alpine location in Sörenberg (Switzerland), we further investigated a pair of sympatric *S. cynipsea* and *S. neocynipsea* populations at high altitude site around Le Mourier in the Cevennes, Southern France (790 m asl), approximately 450 km southwest of the sampling site in Switzerland. Interestingly, we found opposite patterns to those observed at the Sörenberg site (Table 3). Specifically, all tests based on *D*_S_ showed a highly significant excess of ABBA patterns, and hence more interspecific gene flow in sympatry than allopatry. The corresponding tests based on *D*_D_ also indicated an excess of ABBA patterns (recall the difference in sign between *D*_S_ and *D*_D_), but were not significant (Supporting Table S3). In contrast to the patterns in Sörenberg, we did not observe genomic clustering of extreme values of *D*_S_ (Fig. 2, bottom).

**Table 3:** Results of ABBA-BABA tests applied to samples originating from sympatric populations of *S. cynipsea* and *S. neocynipsea* in the French Cevennes as P2 and P3 (or *vice versa*) based on *D*_S_. As in Tables 2 and 3, the upper half of the table summarizes tests involving two *S. neocynipsea* populations from Geschinen (GeN) and Hospental (HoN) as P1 ingroups, and the bottom half summarizes tests involving *S. cynipsea* populations from Pehka, Estonia (PhC) and Petroia, Italy (PtC) as P1.

| P1 | P2 | P3 | $D_S$ | $SD(D_S)$ | $z$ |
| --- | --- | --- | --- | --- | --- |
| **GeN** | **MoN** | **MoC** | **-0.014** | **1.30E-03** | **-11.0** |
| **HoN** | **MoN** | **MoC** | **-0.014** | **1.36E-03** | **-10.6** |
| **PhC** | **MoC** | **MoN** | **-0.030** | **1.84E-03** | **-16.2** |
| **PtC** | **MoC** | **MoN** | **-0.016** | **1.93E-03** | **-8.3** |

## DISCUSSION

We applied the ABBA-BABA approach (Green *et al*., 2010; Durand *et al*., 2011; Soraggi *et al*., 2018) to infer historical gene flow between a pair of closely related species of sepsid flies for which we have detailed information on ecology, morphology, life history, and behavior (Blanckenhorn, 1999; Blanckenhorn *et al*., 2000; Eberhard, 1999; Ozerov, 2005; Parker, 1972a,b; Pont & Meier, 2002; Puniamoorthy *et al*., 2009; Rohner, Blanckenhorn & Puniamoorthy, 2016; Ward, 1983; Ward, Hemmi & Rösli, 1992; Giesen *et al*., 2017, 2019). Previous population genetic analyses of microsatellite markers (Baur *et al*., 2020) and hybridization experiments in the laboratory (Giesen *et al*., 2017, 2019) suggested that *S. cynipsea* and *S. neocynipsea* are genetically distinct, but may occasionally hybridize in nature when occurring in sympatry. Here, we tested this hypothesis by analyzing patterns of DNA sequence variation among pools of field-caught and laboratory specimens from multiple sampling sites across Europe at which either both species occur in sympatry or only one species (*S. cynipsea*) occurs. As discussed in detail below, depending on the focal site of sympatry, we found two qualitatively opposite patterns of interspecific gene flow in sympatry versus allopatry: a relative reduction of gene flow in sympatry at two sampling sites in Switzerland, and a relative excess of gene flow in sympatry at a sampling site in Southern France. Previous studies comparing levels of interspecific gene flow in sympatry and allopatry in other taxa found the full spectrum of results that we here obtained for one species pair. Martin *et al*. (2014; see also Nadeau *et al*., 2013) and Brandvain *et al*. (2014; see also Grossenbacher & Whittall, 2011) found evidence for higher levels of interspecific gene flow in sympatry than in allopatry when studying *Heliconius* butterflies and monkey flowers (*Mimulus guttatus/nasutus*), respectively. In contrast, less gene flow in sympatry than allopatry was observed in studies on *Drosophila arizonae/mojavensis* (Massie & Makow, 2005) and on two species of sea squirt (Tunicata: Bouchemousse *et al*., 2016), while no geographic variation in levels of gene flow between sympatric and allopatric populations was found in studies on wild tomatoes (Nakazato *et al*., 2010). A reduction in average gene flow in sympatry compared to allopatry may suggest reinforcement by natural or sexual selection (Butlin, 1995; Noor, 1999; Coyne & Orr, 2004; cf. Giesen *et al*., 2017, 2019). However, we caution that comparing average levels of gene flow between sympatric and allopatric population pairs may fall short of detecting selection against interspecific gene flow. In monkey flowers, there is evidence for divergent selection against gene flow in sympatry (in terms of a negative correlation between nucleotide divergence and recombination rate) even though average levels of introgression are higher in sympatry than allopatry (Brandvain *et al*. 2014; Aeschbacher *et al*. 2017).

Similar to research on *Heliconius* (Martin *et al*., 2014), we found evidence for an excess of average levels of interspecific gene flow in sympatry compared to allopatry in flies from Le Mourier based on pool-sequenced field-caught specimens (Table 3; Table S3). In contrast, at two Swiss sites where the two species also occur in sympatry, we observed reduced average levels of interspecific gene flow relative to interspecific pairs of parapatric and allopatric populations, similar to patterns found in *Drosophila arizonae/mojavensis* (Massie & Makow, 2005) and sea squirts (Bouchemousse *et al*., 2016). Consistent with our previous study showing behavioral reinforcement after enforced hybridization in the laboratory (Giesen *et al*., 2017), the genomic data analyzed here suggest that selection against interspecific gene flow (Noor, 1999) might also occur in nature in the two sympatric Swiss populations. The reduction of interspecific gene flow was more pronounced at the Sörenberg site in the Swiss Alps (Table 2; Table S1) than at the lowland site in Zürich (Tables 3; Table S2), but qualitatively consistent. However, as evidenced by our contrasting findings for the site in Le Mourier, reinforcement may not be a necessary outcome whenever *S. cynipsea* and *S. neocynipsea* occur in sympatry. The extent of gene flow between these two dung fly sister species thus appears to covary with local environmental conditions. Well-documented differences in climate between Southern France and the Swiss Alps, and between the Swiss Alps and the Swiss lowlands, might provide an ecological explanation for the qualitative and quantitative differences in interspecific gene flow we observed at the different locations (Doebeli & Dieckmann, 2003; Rohner *et al*., 2015). While average minimum temperatures are similar in the French Cevennes and the Swiss Alps, average maximum temperatures are up to 5 °C higher at the French site compared to Sörenberg (Figure 3). In addition, both sampling sites are characterized by very different precipitation regimes. During the main growing season from May until October, Sörenberg (a peat bog site) has on average twice as much rainfall than the location in Le Mourier. However, the mechanisms by which climate affects behavior and/or fitness remain to be understood (Giesen *et al*., 2017, 2019).

**Fig. 3.**
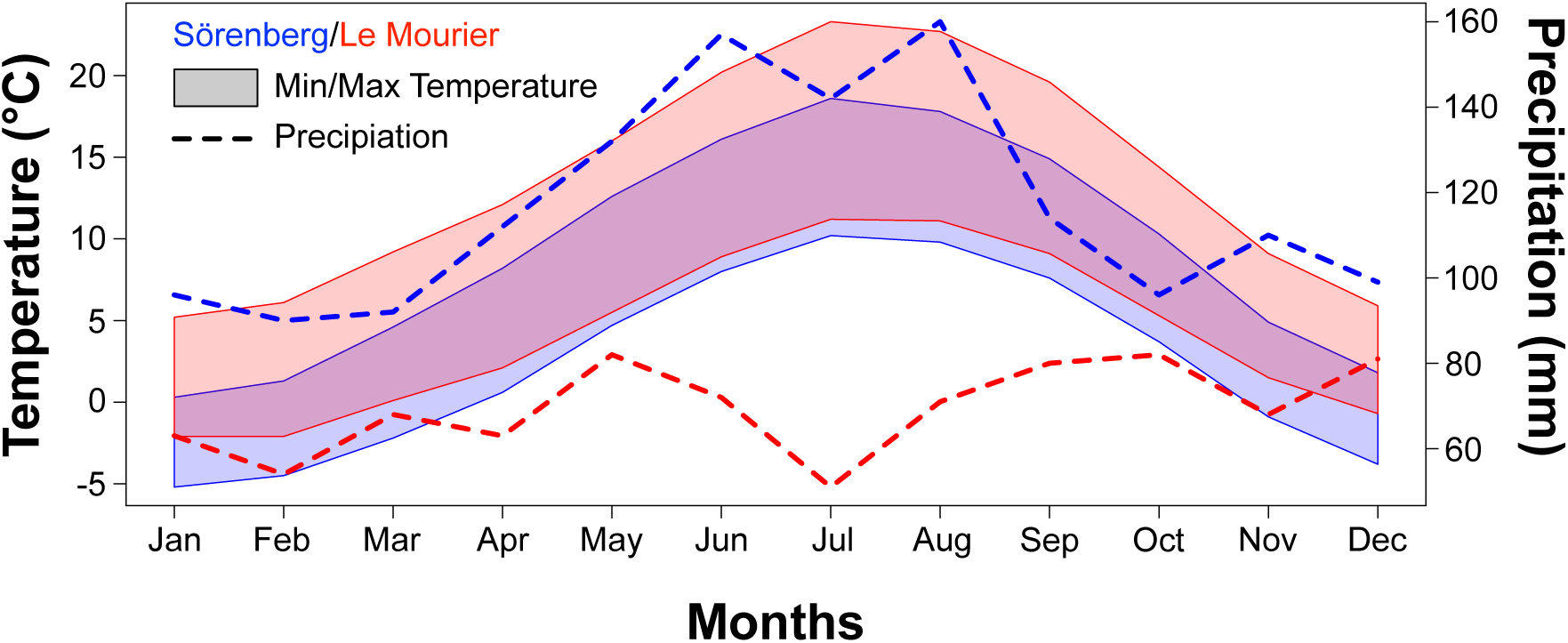
Climatic differences between the Sörenberg (Swiss Alps; blue) and Le Mourier (French Cevennes; red); polygons and dashed lines show monthly average temperature ranges and precipitation at the two sampling sites, respectively. The data from the WorldClim dataset (Hijmans *et al*. 2011) represent 50-year averages of observations in quadratic grid cells with and edge of length 2.5’ (∼ 5 km^2^ in size) around the coordinates of the two sites (Table1). The two locations are characterized by strong differences in precipitation, especially during the main reproductive season of sepsid flies from May to October.

We found a weaker signal of reinforcement between *S. cynipsea* and *S. neocynipsea* at the lowland Zürich site than at the high altitude Sörenberg site in the Swiss Alps. Differences in the strength of reinforcement may also result from climatic variation between these two sites in Switzerland to ultimately influence the strength of selection on natural hybrids. While precipitation patterns of both locations are comparable, mean temperature is approximately 4 °C lower in Sörenberg than in Zürich (Supplementary Figure 1). Alternatively, differences between the two Swiss sites might relate to *S. neocynipsea* being rare around Zürich, in fact in European lowlands in general (Pont & Meier, 2002), while this species is common at higher alpine altitudes. If based primarily on behavioral mate choice, reinforcement is expected to be stronger, and hence its evolution more likely, wherever interspecific encounter frequencies are higher. Moreover, given that the evidence for the Zürich site is based on both wild-caught and laboratory specimens, the weaker patterns of reinforcement observed in Zürich might be partially explained by purifying selection in the laboratory against introgressed variants. Nevertheless, the results from our analyses performed with both wild-caught and laboratory *S. cynipsea* samples from Zürich yielded consistent results (Supplementary Table 2), suggesting that whether flies came from a natural or laboratory population did not affect the outcome. Overall, we therefore do not think that purifying selection in the laboratory confounded our results.

Laboratory hybridization experiments have shown that interspecific mating between *S. cynipsea* and *S. neocynipsea* can result in viable and fertile F_1_ hybrid females and offspring in backcrosses with the parental species (Giesen *et al*., 2017, 2019). Together with morphological and behavioral similarities among the two species, these laboratory experiments suggest that hybridization may well take place in areas of co-occurrence, such as the French Cevennes. In other areas, including our sampling site in the Swiss Alps, subtle premating behavioral barriers and the reductions in fecundity and fertility also previously documented in the laboratory by Giesen *et al*. (2017, 2019) may combine with micro-ecological niche differences to mediate spatio-temporally divergent reproductive timing. Such a divergence may effectively prevent hybridization in nature and ultimately reinforce species boundaries. This interpretation is strengthened by a study on Central American *Archisepsis diversiformis* flies showing that mating between two disjunct populations is only evident under forced laboratory conditions, while under conditions of free mate choice flies from the different populations do not interbreed (Puniamoorthy 2014). Behavioral mating barriers therefore seem to evolve comparatively fast (Gleason & Ritchie, 1998; Puniamoorthy *et al*., 2009; Puniamoorthy, 2014), especially if species occur in sympatry and reinforcement by natural or sexual selection can operate (Coyne & Orr, 2004; Seehausen, 2004; Ritchie, 2007).

Our study of introgression patterns among sympatric *S. cynipsea* and *S. neocynipsea* populations relied on the ABBA-BABA test, which was initially designed for the analysis of haploid genotypes of one individual from each of four lineages (Green *et al*., 2010). Recent extensions of this original approach to allele frequency data by Durand *et al*. (2011) and Soraggi *et al*. (2018), which incidentally were here shown to not be equally sensitive in picking up introgression patterns, enabled us to apply the method equivalently to pool-sequenced laboratory as well as to field-caught specimens (see also Deitz *et al*. 2016). As a technical innovation, we have here provided a Python implementation of the ABBA-BABA approaches by Durand *et al*. (2011) and Soraggi *et al*. (2018) that takes a simple 2D matrix of allele frequencies as input. This implementation allows restricting the ABBA-BABA test to high-confidence SNPs that pass a set of user-defined filters. Such filtering can reduce artefacts of pool-sequencing (Schlötterer *et al*. 2014) that may produce false polymorphisms due to sequencing errors (Futschik & Schlötterer 2011; Cutler & Jensen 2012). In addition, and in contrast to the software package ANGSD (Korneliussen *et al*. 2014), our implementation accommodates an outgroup (P4) taxon different from the one used as a reference to call the SNPs. This feature reduces a potential reference bias if the ingroup taxa (P1, P2, P3) strongly differ from an outgroup that also serves as reference (Ballouz *et al*. 2019). Our script and detailed instructions for how to use it are available on GitHub (https://github.com/capoony/ABBA-BABA-4-AF; see also Supporting Information).

In summary, we found evidence that the extent of interspecific gene flow between two closely related sepsid fly species in sympatry relative to allopatry varies with the geographic location of the site. Interspecific gene flow in sympatry appears to be reduced at two Swiss sites, where the two species have likely locally adapted to different (micro-)ecological niches, such as different breeding times or phenologies, and/or where pre- or postmating reproductive barriers (Giesen *et al*., 2017) have been and are subject to stronger reinforcement. Future studies are needed to determine if ecological and/or sexual selection are indeed driving the gene flow patterns we identified, and future genomic research should analyze patterns of genomic differentiation across allopatric populations from Europe as well as North America (Baur *et al*., 2020). It remains unclear why *S. neocynipsea* is rare in Europe, but common in North America, and why *S. cynipsea* abounds around fresh dung all over Europe north of the Alps and apparently outcompetes and ultimately relegates *S. neocynipsea* towards presumably marginal, high altitude habitats (Pont & Meier, 2002). Such analyses would provide further insights into introgression patterns across the species’ entire natural range, as well as into the role of local (e.g., longitudinal or latitudinal) adaptation in shaping genome-wide patterns of differentiation.

## Supporting information

Supplementary Figures Tables and additional analyses

## ACKNOWLEDGEMENTS

Genome resequencing was performed at the Functional Genomics Center Zürich (FGCZ), and we particularly thank FGCZ staff member Jelena Kühn-Georgijevic for her support. We thank the Wolfpack at the University of Zürich for their support and help in maintaining fly cultures, and Patrick Rohner, Juan Pablo Busso, Nalini Puniamoorthy, Anders Kjaersgaard, Toomas Esperk and Cait Dmitriew for contributing samples. B.M., O.N. and L.P. thank the Leibniz association for granting funds to establish the Leibniz graduate school for Genomic Biodiversity Research (GBR), in which context the reference draft genomes of several *Sepsis* species explored in this study were sequenced and assembled. We particularly thank the GBR members Jeanne Wilbrandt, Jan Philip Øyen, and Malte Petersen for their contribution to the assembly pipeline. The University Research Priority Program ‘Evolution in Action’ of the University of Zürich, the Swiss National Science Foundation, and the Georges and Antoine Claraz-Donation funded the work of A.G., the Austrian Science Foundation (grant no. FWF P32275) the work of M.K.

## DATA ACCESSIBILITY

All sequencing data have been deposited at the short-read archive (SRA; https://www.ncbi.nlm.nih.gov/sra) under the accession number PRJNA612154. Novel software is available at GitHub (https://github.com/capoony/ABBA-BABA-4-AF).

## AUTHORS CONTRIBUTION

AG, WUB, KKS, HELL, SA and MK conceived and designed the study. MAS and AG gathered and prepared the samples and generated the genomic data. BM, ON and LP sequenced and constructed the reference genome. MK, AG, SA and HELL analyzed the genomic data. MK developed new software. AG, MK, SA and WUB wrote the manuscript, with ON, KKS, RSI, LP, BM, MAS and HELL contributing to the interpretation of the data, and editing and commenting on various versions of the manuscript. WUB, KKS, RSI, and HELL were members of AG’s PhD committee, providing guidance and input at all stages.

