## Supplementary Figures Tables and additional analyses for "Genomic signals of admixture and reinforcement between two closely related species of European sepsid flies"

**Supplementary Material**

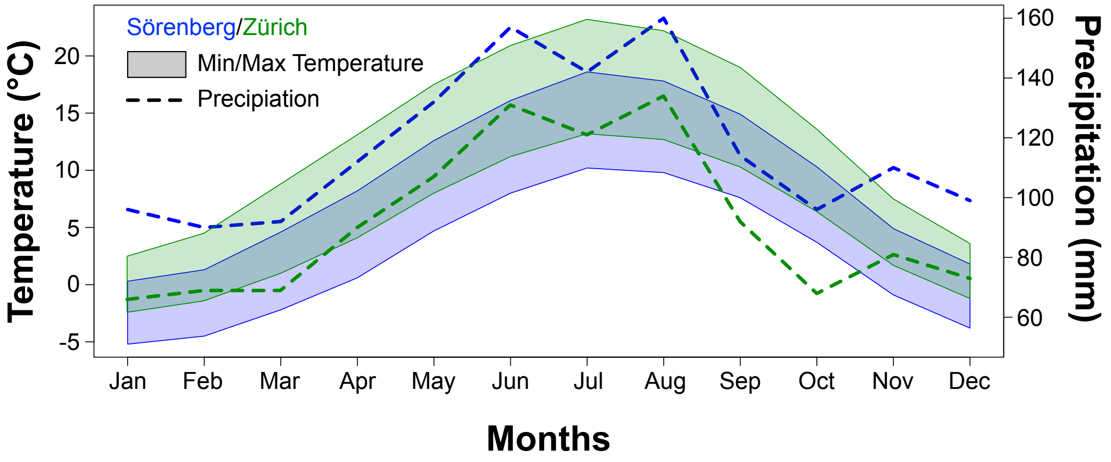


**Supplementary Figure S1:** Polygons and dashed lines show monthly average temperature ranges and precipitation at the two sampling sites in Sörenberg (Switzerland, blue) and Zürich (Switzerland, red), respectively. The data from the WorldClim dataset (Hijmans *et al*. 2011) represent 50-year averages of observations in quadratic grid cells with an edge of length 2.5’ (~ 5 $km^2$ in size) around the corresponding coordinates (Table 1).

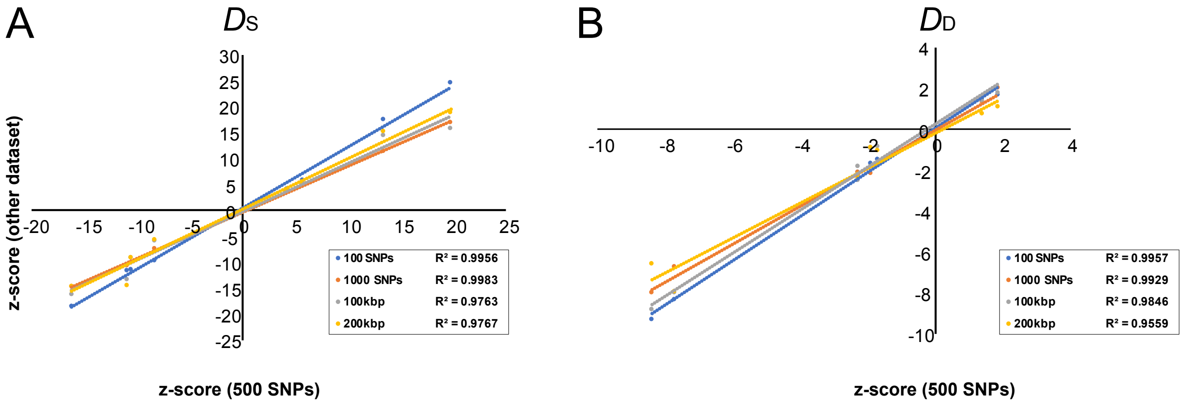


**Supplementary Figure S2:** Correlations between the *z*-scores of our chosen ABBA-BABA reference window size of 500 consecutive SNPs on the *x* axis with those of various alternative genomic window sizes (100 or 1000 consecutive SNPs, 100 or 200 kb) on the *y* axis. Each (differently colored) comparison entails the *z*-scores from the 6 possible ABBA-BABA tests involving the Sörenberg and Le Mourier sympatric populations as P2 and P3, and PhC, PtC, GeN and HoN populations as P1. The coefficients of determination ($R^2$) indicate how much of the variance is explained by the corresponding linear regression in *R*. Panels A and B show results based on $D_S$ (Soraggi *et al*. 2018) and $D_D$ (Durand *et al*. 2011), respectively.

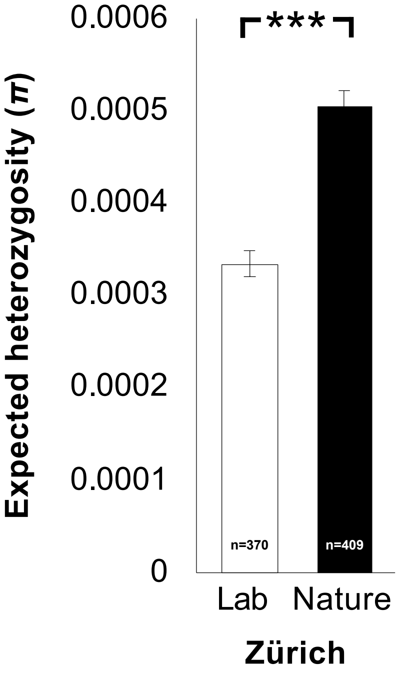


**Supplementary Figure S3:** Expected heterozygosity ($\pi$) per SNP in laboratory populations (after at least 30 generations of breeding) and field-caught samples of *S. cynipsea* from the Zürich site. Expected heterozygosity was estimated in non-overlapping sliding windows of 200 kb along the genome using corrections for pooled resequencing (Futschik & Schlötterer 2010) as implemented in *PoolGen* (https://github.com/capoony/DrosEU_pipeline; Kapun *et al*. 2018). Error bars represent standard errors. The number of 200-kb windows analyzed are shown at the bottom of each bar. The difference in heterozygosity is significant (Mann Whitney U test, $U = 55558$, $p < 0.001$)

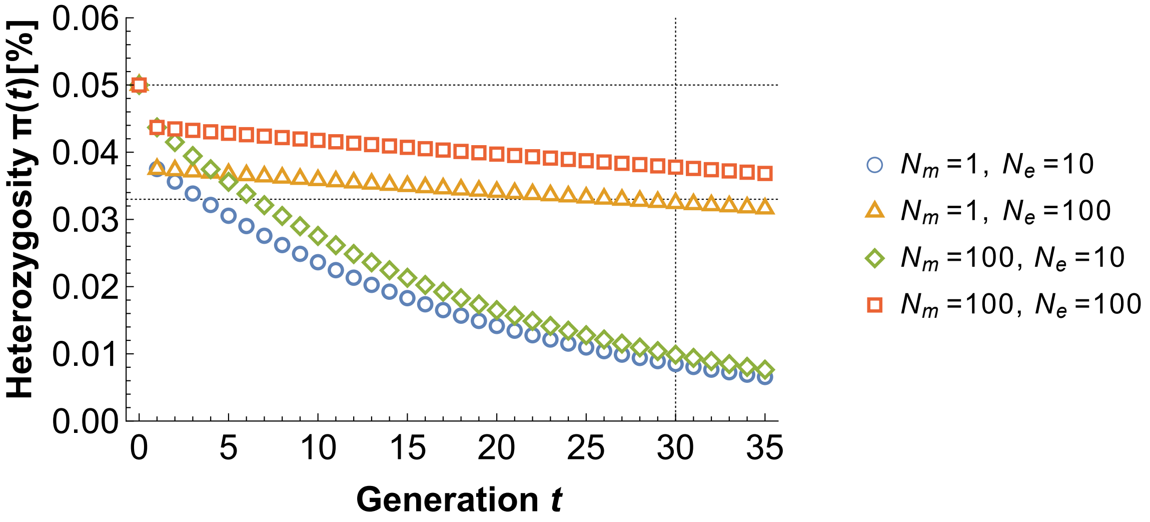


**Supplementary Figure S4:** Dynamics of the expected heterozygosity $\pi$ after a founder event and subsequent breeding in the laboratory according to Eq. S1. The dynamics are shown for a single female *S. cynipsea* fly that had mated with $N_m$ males in nature prior to being used to initiate a laboratory population of constant effective size $N_e$. Horizontal black lines indicate the estimated expected heterozygosity in the natural *S. cynipsea* population at the Zürich site [π(0), top line] and in the derived laboratory population [π(30), bottom line] at generation $t = 30$ (vertical black line). Values of π(0) and π(30) corresponding to the two horizontal dashed lines are taken from Supplementary Figure S3. Combinations of $N_m$ and $N_e$ yielding dynamics of π that intersects with the lower black line at generation 30 can plausibly explain the observed data. Here, this is the case for the parameter combination represented by the orange dots ($N_m = 1, N_e = 100$), but not for the other parameter combinations.

**Supplementary Table S1:** Results of ABBA-BABA tests applied to samples originating from sympatric populations of *S. cynipsea* and *S. neocynipsea* in Sörenberg as P2 and P3 (or *vice versa*) based on the $D_D$-statistics (Durand *et al.* 2011). The upper half of the table shows tests with two European *S. neocynipsea* populations from Geschinen (GeN) and Hospental (HoN) as P1 ingroups, and the bottom half shows the reciprocal approach with two Swiss *S. cynipsea* populations from Pehka, Estonia (PhC) and Petroia, Italy (PtC) as P1 ingroups. The first three columns indicate the phylogeny assumed for the test, with P4 always being *S. orthocnemis;* columns 4 to 6 give the genome-wide average of $D_D$, its jackknifed standard deviation SD($D_D$), and the corresponding $z$-score. Significant tests are shown in bold.

| P1 | P2 | P3 | $D_D$ | SD($D_D$) | $z$ |
|---|---|---|---|---|---|
| GeN | SoN | SoC | 0.007 | 5.09E–03 | 1.4 |
| HoN | SoN | SoC | 0.010 | 5.27E–03 | 1.9 |
| **PhC** | **SoC** | **SoN** | **–0.052** | **6.19E–03** | **–8.4** |
| **PtC** | **SoC** | **SoN** | **–0.049** | **6.34E–03** | **–7.7** |

**Supplementary Table S2:** Results of ABBA-BABA tests applied to samples originating from sympatric populations of *S. cynipsea* and *S. neocynipsea* in Zürich as P2 and P3 (and vice versa) based on $D_S$ (top table above double-line) and $D_D$ (bottom table below double-line). As in Table 2, the upper halfs of each table summarize tests involving two *S. neocynipsea* populations from Geschinen (GeN) and Hospental (HoN) as P1 ingroups, and the bottom half summarizes tests involving *S. cynipsea* populations from Pehka, Estonia (PhC) and Petroia, Italy (PtC) as P1. Other details as in Table 2. The first three columns indicate the phylogeny assumed for the test, with P4 always being *S. orthocnemis*; columns 4 to 6 give the genome-wide average of $D_S$ and $D_D$, its jackknifed standard deviation SD($D_S$) and SD($D_D$), and the corresponding $z$-score. Significant tests are shown in bold.

| P1 | P2 | P3 | $D_S$ | SD($D_S$) | $z$ |
|---|---|---|---|---|---|
| **GeN** | **IZuN** | **IZuC** | **0.016** | **2.63E–03** | **6.1** |
| **GeN** | **IZuN** | **ZuC** | **0.055** | **2.79E–03** | **19.6** |
| **HoN** | **IZuN** | **IZuC** | **0.017** | **2.68E–03** | **6.5** |
| **HoN** | **IZuN** | **ZuC** | **0.055** | **2.69E–03** | **20.4** |
| PhC | IZuC | IZuN | 0.001 | 3.89E–03 | 0.3 |
| PhC | ZuC | IZuN | –0.002 | 1.54E–03 | –1.4 |
| **PtC** | **IZuC** | **IZuN** | **0.012** | **3.74E–03** | **3.3** |
| **PtC** | **ZuC** | **IZuN** | **0.009** | **1.66E–03** | **5.3** |

| P1 | P2 | P3 | $D_D$ | SD($D_D$) | $z$ |
|---|---|---|---|---|---|
| **GeN** | **IZuN** | **IZuC** | **0.030** | **9.35E–03** | **3.2** |
| GeN | IZuN | ZuC | 0.006 | 8.88E–03 | 0.7 |
| HoN | IZuN | IZuC | 0.024 | 9.41E–03 | 2.5 |
| HoN | IZuN | ZuC | 0.005 | 9.33E–03 | 0.6 |
| PhC | IZuC | IZuN | –0.014 | 9.77E–03 | –1.4 |
| **PhC** | **ZuC** | **IZuN** | **–0.019** | **5.61E–03** | **–3.4** |
| PtC | IZuC | IZuN | –0.012 | 9.91E–03 | –1.2 |
| PtC | ZuC | IZuN | –0.016 | 5.94E–03 | –2.7 |

**Supplementary Table S3**: Results of ABBA−BABA tests applied to samples originating from sympatric populations of *S. cynipsea* and *S. neocynipsea* in the French Cevennes as P2 and P3 (and *vice versa*) based on $D_D$. As in Supplementary Table S1, the upper half of the table summarizes tests involving two *S. neocynipsea* populations from Geschinen (GeN) and Hospental (HoN) as P1 ingroups, and the bottom half summarizes tests involving *S. cynipsea* populations from Pehka, Estonia (PhC) and Petroia, Italy (PtC) as P1.

| P1 | P2 | P3 | $D_D$ | SD($D_D$) | $z$ |
|---|---|---|---|---|---|
| GeN | MoN | MoC | −0.011 | 5.03E−03 | −2.3 |
| HoN | MoN | MoC | −0.012 | 5.14E−03 | −2.3 |
| PhC | MoC | MoN | −0.010 | 5.37E−03 | −1.9 |
| PtC | MoC | MoN | −0.009 | 5.24E−03 | −1.7 |

**Overview of ABBA-BABA test statistics**

In the original ABBA-BABA test, Green *et al.* (2010) computed the *D*-statistic across *n* bases as

$$D_{\mathrm{G}} = \frac{\sum_{i=1}^{n}[C_{\mathrm{ABBA}}(i) - C_{\mathrm{BABA}}(i)]}{\sum_{i=1}^{n}[C_{\mathrm{ABBA}}(i) + C_{\mathrm{BABA}}(i)]},$$

where $C_{\mathrm{ABBBA}}(i)$ and $C_{\mathrm{BABA}}(i)$ are indicator variables taking a value of 0 or 1 depending on whether an ABBA or BABA configuration is observed at base $i$. This version of the test was developed for aligned sequence data from four haploid individuals (or, equivalently, diploid individuals made pseudo-haploid by random sampling of alleles). To extend the scope of the ABBA-BABA test to allele frequencies at a set of SNPs, Durand *et al.* (2011) proposed the modified *D* statistic

$$D_{\mathrm{D}} = \frac{\sum_{i=1}^{n}[(1-\hat{p}_{i1})\hat{p}_{i2}\hat{p}_{i3}(1-\hat{p}_{i4}) - \hat{p}_{i1}(1-\hat{p}_{i2})\hat{p}_{i3}(1-\hat{p}_{i4})]}{\sum_{i=1}^{n}[(1-\hat{p}_{i1})\hat{p}_{i2}\hat{p}_{i3}(1-\hat{p}_{i4}) + \hat{p}_{i1}(1-\hat{p}_{i2})\hat{p}_{i3}(1-\hat{p}_{i4})]},$$

where $\hat{p}_{ij}$ is the estimated frequency of the derived allele *B* at SNP *i* in population (species) $j$. An excess of the ABBA allele pattern thus results in a positive $D_G$ $(D_D)$, whereas an excess of the BABA allele pattern results in a negative $D_G$ $(D_D)$.

Recently, Soraggi *et al.* (2018) proposed an extension of the original ABBA-BABA test that accommodates data at *M* genomic sites from multiple individuals per population sequenced at varying depth. Soraggi *et al.* (2018) expressed the probabilities of observing ABBA and BABA patterns in terms of their expected values with respect to the population allele-frequency distributions,

$$\Pr(\mathrm{ABBA}_i) = E[(1-p_{i1})p_{i2}p_{i3}(1-p_{i4}) + p_{i1}(1-p_{i2})(1-p_{i3})p_{i4}],$$

and

$$\Pr(\mathrm{BABA}_i) = E[(1-p_{i1})p_{i2}(1-p_{i3})p_{i4} + p_{i1}(1-p_{i2})p_{i3}(1-p_{i4})].$$

Under the null hypothesis of no gene flow, the probabilities $\Pr(\mathrm{ABBA}_i)$ and $\Pr(\mathrm{BABA}_i)$ are equal, i.e.

$$H_0\colon \Pr(\mathrm{BABA}_i) - \Pr(\mathrm{ABBA}_i) = E[(p_{i1}-p_{i2})(p_{i3}-p_{i4})] = 0 \text{ for all } i = 1, \ldots, M.$$

Equivalent to the approach of Durand *et al.* (2011), Soraggi *et al.* (2018) normalized by

$$\Pr(\mathrm{BABA}_i) + \Pr(\mathrm{ABBA}_i) = E[(p_{i1}+p_{i2}-2p_{i1}p_{i2})(p_{i3}+p_{i4}-2p_{i3}p_{i4})].$$

For empirical allele frequencies $\hat{p}_{ij}$ as unbiased estimators of $p_{ij}$ $(j = 1, \ldots, 4)$, Soraggi *et al.* (2018) defined their extended *D*-statistic as

$$D_{\mathrm{S}} = \frac{\sum_{i=1}^{n}[(\hat{p}_{i1}-\hat{p}_{i2})(\hat{p}_{i3}-\hat{p}_{i4})]}{\sum_{i=1}^{n}[(\hat{p}_{i1}+\hat{p}_{i2}-2\hat{p}_{i1}\hat{p}_{i2})(\hat{p}_{i3}+\hat{p}_{i4}-2\hat{p}_{i3}\hat{p}_{i4})]},$$

and showed that the distribution of $D_{\mathrm{S}}$ converges to a Gaussian distribution under $H_0$. Note that, due to how $H_0$ was formulated, $D_{\mathrm{S}}$ has the opposite sign of $D_{\mathrm{D}}$ and $D_{\mathrm{G}}$, i.e. and excess (deficiency) of the BABA (ABBA) allele pattern implies $D_{\mathrm{S}} > 0$, and *vice versa*. We followed

Soraggi *et al*. (2018) in restricting our analyses to informative sites only, i.e. sites at which samples from P1 and P2, or P3 and P4 (cf. above and Fig. 1) are not fixed for the same allele.

### Implementation of ABBA-BABA tests based on allele frequency data

The two approaches to calculating *D*-statistics by Green *et al*. (2010) and Soraggi *et al*. (2018) are implemented in the software ANGSD (Korneliussen, Albrechtsen & Nielsen, 2014). While ANGSD integrates over site-specific genotype likelihoods when calculating *D*, it currently does not support pool-sequencing data (but see Deitz *et al*. 2016). Moreover, ANGSD does not provide the possibility to use a pre-defined set of high-confidence SNPs, since variants are called based on the provided samples as part of the analysis. We therefore implemented the calculation of the *D*-statistics from allele frequency data by Durand *et al*. (2011) and by Soraggi *et al*. (2018) (Python script available from https://github.com/capoony/ABBABABA-4AF). Our script calculates *D* both across the entire genome as well as within non-overlapping genomic windows defined to all either contain the same number of SNPs or to be equal in sequence length. To test for significant deviations of *D* from 0, our script also calculates *z*-scores based on jackknifing following the approach suggested by Busing *et al*. (1999). When using windows with equal numbers of SNPs, we perform a blocked, even *m*-delete jackknife procedure, where *m* is the group size of observations removed from the sample for jackknifing. This differs from the method implemented in ANGSD. ANGSD uses an uneven *m*-delete procedure because it uses windows of equal sequence length, which may contain unequal numbers of SNPs. We removed one window (block) at a time to obtain standard deviations and calculate *z*-scores following Green *et al*. (2010). We adopted the commonly used significance threshold of $|z| > 3$ (Reich *et al*., 2011; Jeong *et al*., 2016; Novikova *et al*., 2016). In addition, our implementation also permits estimating variances by a blocked, uneven *m*-delete jackknife procedure similar to ANGSD whenever block-wise *D* is calculated in windows based on sequence length.

In contrast to Soraggi et al. (2018), who computed the *D*-statistic for human genomic data in windows of 5 million base pairs, we chose equisized blocks of 500 SNPs for the calculation of *D* given that sepid genomes are ~20-fold smaller than the human genome. Moreover, keeping the number of SNPs constant in each block results in similar variances across blocks and facilitates the estimation of variances using the even *m*-block jackknife method (Busing et al. 1999). However, to test for the robustness of our approach, we repeated the analyses with different block sizes (based on 100, 500 and 100 SNPs) and sequence length (100000 and 200000 bp) and found highly significant correlations among the *z*-scores of the different analyses (see Supplementary Figure S2). This result indicates that our ABBA-BABA results are not influenced by the choice of window-size and the type of blocking.

**Reduction of genetic diversity in laboratory populations**

We expected the genetic diversity in the laboratory population of *S. cynipsea* to be strongly reduced relative to that of recently field-caught flies from the same (Zürich) site. While we did observe significantly lower levels of expected heterozygosity (Nei 1987) in laboratory than field-caught populations (Mann Whitney U test, $p < 0.001$; Supplementary Figure S3), reductions in heterozygosity were much lower than expected under strong inbreeding. Our laboratory populations were therefore not highly inbred, so we decided to treat the sequencing data from laboratory and natural populations equally in all our analyses.

To interpret the decrease in genetic diversity that we observed in our laboratory populations relative to field-caught flies we adopted a simple population genetic model for the decay of the expected heterozygosity $\pi$ (Nei 1987). Specifically, we assumed that a single wild-caught female mated to $N_m$ males was brought into the laboratory to initiate a panmictic population of effective size $N_e$, and that this population was then maintained for $n$ generations. According to Eq. (3.122) of Ewens (2004, p. 123), and given a single founding female, we approximated the effective population size during the founder event at generation 0 by $4N_m/(1 + N_m)$. The expected heterozygosity at generation $t$ is then given by

$$\pi(t) = \left(1 - \frac{1}{2\times 4N_m/(1+N_m)}\right)\left(1 - \frac{1}{2N_e}\right)^{t-1}\pi_0, \qquad \text{(Eq. S1)}$$

where $\pi_0$ is the expected heterozygosity in the natural population. By substituting the expected heterozygosity computed from allele frequencies in field-caught and laboratory populations after $n = 30$ generations for $\pi_0$ and $\pi(t = 30)$, respectively, we explored which combinations of $N_m$ and $N_e$ would explain the observed reduction in $\pi$ (Supplementary Figure S1). We limited $N_e$ to 100, which is our upper estimate for the census size of the laboratory populations. We set $\pi_0 = 0.05$ % and $\pi(30) = 0.033$ % according to our estimates. This 34 % reduction in heterozygosity is compatible with a founding female mated to a single male ($N_m = 1$) and a subsequent laboratory effective population size as large as $N_e = 100$ (Supplementary Figure S4). These results suggest that the initial founder event and successive breeding in the laboratory did not induce strong inbreeding.
